# Hypoxia-mediated regulation of mitochondrial transcription factors: Implications for hypertensive renal physiology

**DOI:** 10.1101/816470

**Authors:** Bhargavi Natarajan, Vikas Arige, Abrar A. Khan, S. Santosh Reddy, Rashmi Santhoshkumar, B. K. Chandrasekhar Sagar, Manoj K. Barthwal, Nitish R. Mahapatra

## Abstract

Kidneys have a high resting metabolic rate and low tissue partial pressure of oxygen due to enhanced mitochondrial oxygen consumption and ATP production for active solute transport. Enhanced mitochondrial activity leads to progressive hypoxia from the renal cortex to renal medulla. Renal tubulointerstitial hypoxia (TiH) is severe in hypertensive rats due to increased sodium reabsorption within their nephrons. Additionally, these rats display increased energy demand and therefore, require healthy mitochondria for adequate salt reabsorption. Hence, we sought to study the regulation of mitochondrial biogenesis and expression of mitochondrial transcription factors (mtTFs, viz. Tfam, Tfb1m and Tfb2m) during hypoxic conditions and in rodent models of genetic hypertension. We report that the expressions of HIF-1α (hypoxia inducible factor-1α), PGC-1α (peroxisome proliferator activated receptor-γ co-activator-1α), mtTFs and OXPHOS proteins are elevated in hypertensive rats as compared to their normotensive counterparts. Additionally, studies in cultured kidney cells show that acute hypoxia augments the expression of these genes. We also observe a positive correlation between HIF-1α and mtTFs transcripts in human tissues. Furthermore, we report for the first time to our knowledge, that HIF-1α binds to promoters of Tfam, Tfb1m and Tfb2m genes and augments their promoter activities in NRK52e cells subjected to acute hypoxia. Taken together, this study suggests that acute hypoxia may enhance mitochondrial function to meet the energy demand in renal tubular epithelial cells and in young/pre-hypertensive SHR kidneys.

**Translational Statement:** Our results suggest that tubulointerstitial hypoxia (TiH) prevailing in prehypertensive rats augments the expression of mitochondrial transcription factors and proteins of electron transport chain. Moreover, previous reports indicate that ATP synthesis in these rats are elevated. Thus, our study provides insights into the molecular mechanism of such enhanced mitochondrial function. We propose that during early stages of kidney diseases (marked by mild TiH) an enhancement of mitochondrial function via stimulation of HIF-1α/PGC-1α production may delay renal tubular damage.

## INTRODUCTION

Kidneys play a key role in blood pressure regulation. Active solute transport by kidneys increases oxygen consumption and thereby, causes hypoxic environment within the renal epithelial cells, termed as tubulointerstitial hypoxia (TiH). Intracellular hypoxia then triggers reprogramming of cellular activities to meet the energy requirement^1, 2^. Therefore, it has been hypothesized that kidneys operate at the brink of anoxia^3, 4^.

Multiple studies demonstrate renal mitochondrial dysfunction in hypertensive rodent models viz. Spontaneously Hypertensive Rats (SHR) and Dahl Salt Sensitive (SS) rats^5–8^. Furthermore, it has been reported kidneys rely majorly on that fatty acid oxidation (FAO) by mitochondria to meet the energy demand^9^. Increased activity of membrane transporters in kidneys of SHR exacerbates TiH which is milder in the normotensive WKY^10^. Defective renal oxygenation and increased mitochondrial oxidative stress have been reported in SHR^11, 12^. Oxygen consumption rate and ATP synthesis were found to be higher in the proximal tubular cells of young SHR when compared with age-matched WKY^13–15^. Kidney transplantation studies provide evidence that renal genetic factors may be responsible for hypertension progression in SHR^16–19^. However, the renal expression of mitochondrial genes that could explain TiH and enhanced ATP production in SHR is largely elusive. Specifically, the expression of mitochondrial transcription factors under hypertensive and hypoxic conditions remains unexplored.

Mitochondrial transcription factors (mtTFs; viz. Tfam, Tfb1m and Tfb2m) are crucial players in mitogenesis and mitochondrial genome expression^20^. Mammalian mitochondrial genome is ∼16 kb and includes 37 genes and a control/regulatory region^21^. Tfam enables both replication and transcription of mitochondrial genome by binding to mitochondrial DNA and RNA polymerases, PolG and Polrmt, respectively. Tfam has also been reported to cooperatively bind to the entire mitochondrial DNA (mtDNA) facilitating mtDNA packaging and thereby, protecting it from ROS-induced damage^22^. Tfb2m binds to Polrmt and enables promoter recognition^23, 24^. Tfb1m aids in translation by initiating mitoribosome biogenesis^25^. Peroxisome-proliferator-activated-receptor-γ-co-activator-1α (PGC-1α), the master regulator of mitogenesis drives the transcription of mtTFs and other mitochondrial genes in association with Nuclear Respiratory Factors 1 and 2 (NRF1/NRF2) and Sp1^26, 27^.

In this study, we investigated the expression of mitochondrial transcription factors and mitochondrial genes in the kidney tissues of SHR and WKY. Furthermore, in order to understand mitochondrial biogenesis in renal hypoxic environment, we used cultured kidney cells to study the regulation of mtTFs and mitochondrial genes under acute hypoxia. Interestingly, we found that mtTFs and OXPHOS proteins are elevated both in hypoxic hypertensive kidneys of SHR and corroboratively in cultured kidney cells subjected to acute hypoxia.

## MATERIALS AND METHODS

A detailed description of the methods used for this study can be found in the supplementary file.

### Rat strains

Kidney tissues were harvested from young (4-6-week-old) hypertensive SHR and normotensive WKY males. The animal experiments were approved by the Institute Animal Ethics Committee at Indian Institute of Technology Madras. India.

### Generation of mtTFs promoter-reporter constructs

The mtTFs promoter-reporter constructs were generated in the pGL3Basic vector (Promega, USA). The resultant plasmids were sequenced for confirmation. Primers used for cloning are listed in Table 1.

**Table 1:**
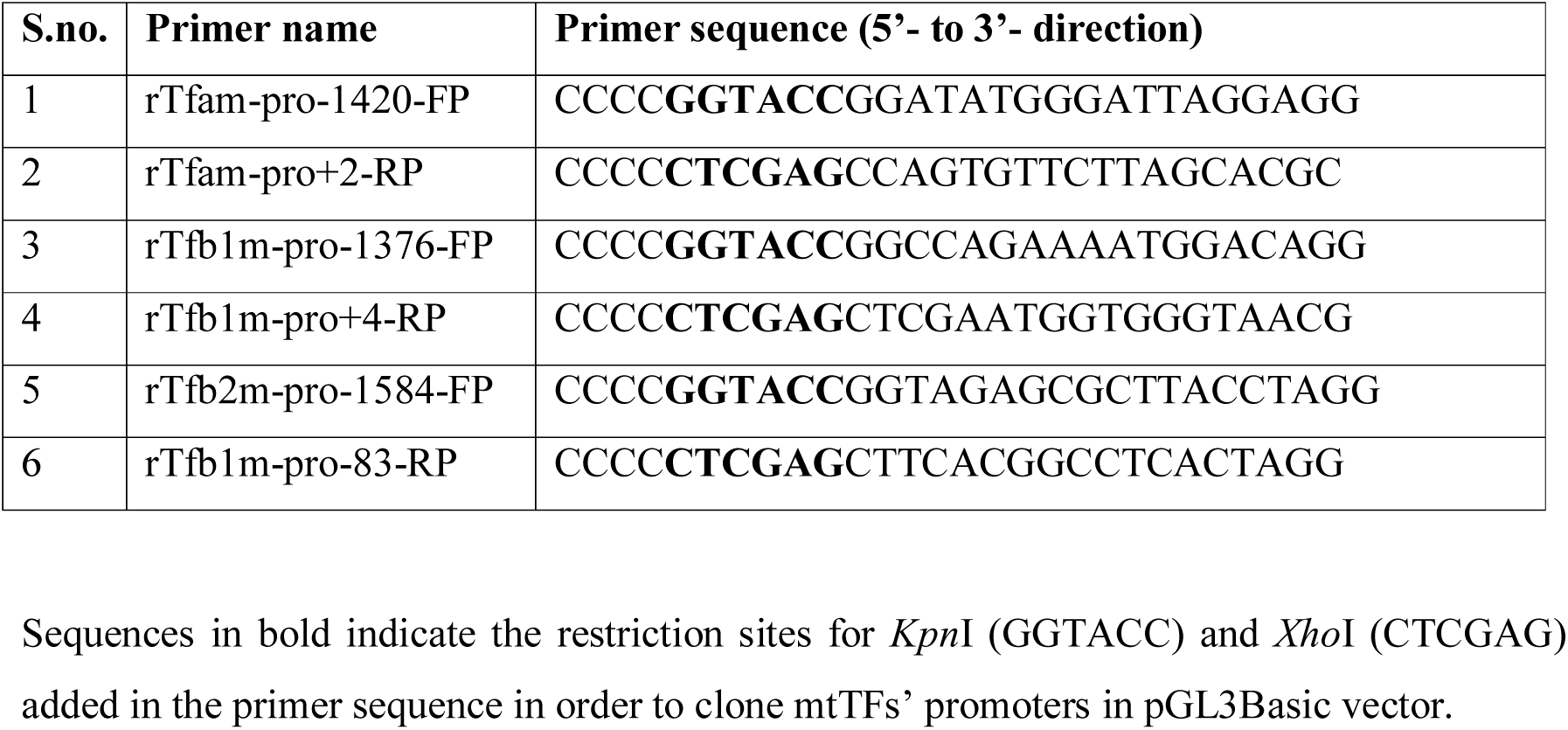
Primers for cloning the promoter region of mitochondrial transcription factors.

### *In silico* prediction of Hypoxia Response Elements in mtTFs’ promoters and their conservation across mammals

Online transcription factor binding site tools (viz. ConSite, Matinspector, LASAGNA, Jaspar and p-Match) were used to predict potential Hypoxia Response Elements (HREs) in the promoters of mtTFs. Genedoc software was utilized to analyze the conservation of HREs among various mammals.

### Cell culture, transfections, hypoxia and reporter assays

Authenticated Normal Rat Kidney epithelial cells (NRK 52e) were cultured and transfected with 500 ng of the desired promoter-reporter constructs. Acute hypoxia was established using 99% argon gas and luciferase activity was determined.

### DNA/RNA isolation and quantitative real-time PCR

The relative expression of mtTFs transcripts, mitochondrial RNA (mtRNA) and mitochondrial DNA (mtDNA) levels in animals tissues and cultured cells were analyzed through qPCR using the ΔΔCt method as described previously^28, 29^. The primers used are listed in Table 2.

**Table 2:**
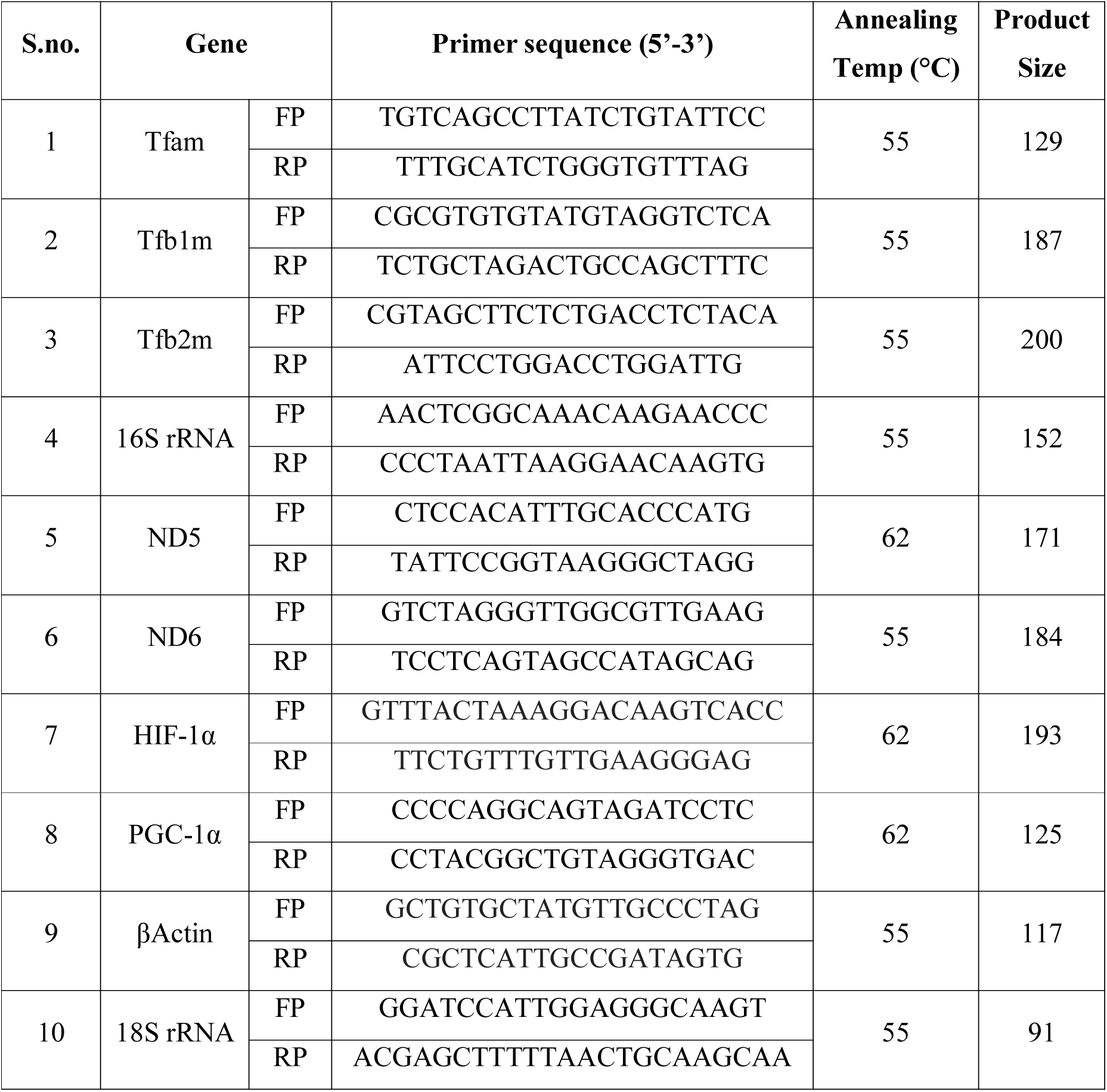
Primers used for qPCR to quantify various mitochondrial genes.

### Western blotting for various proteins in NRK52e cells and rat model of essential hypertension

The protein expression of mtTFs and other mitochondrial proteins were determined by Western blot analysis using specific antibodies.

### Chromatin immunoprecipitation

Chromatin immunoprecipitation assays were using our method described previously^28^. Purified immuneprecipitated DNA was used for qPCR analysis using primers listed in Table 3.

**Table 3:**
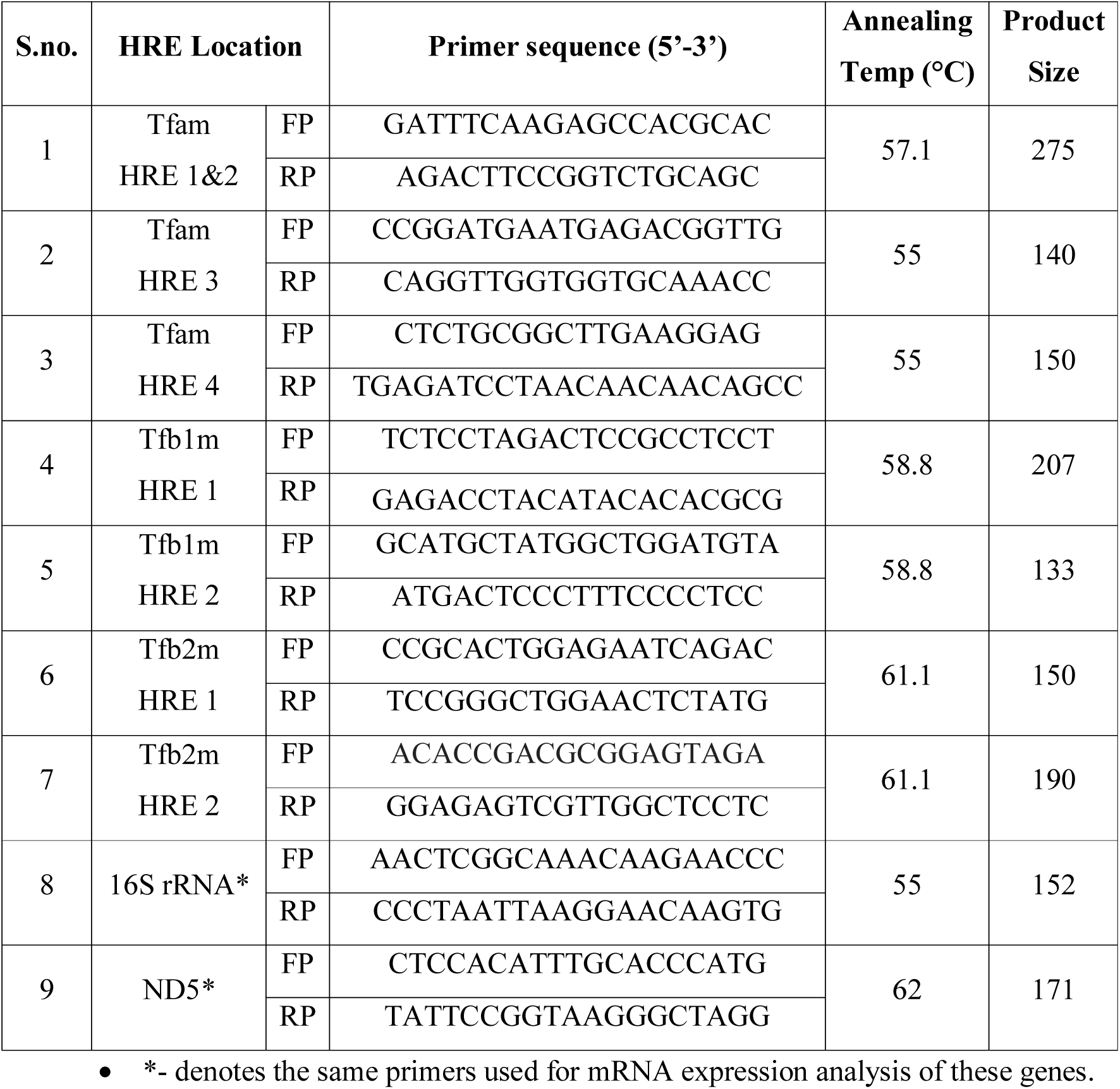
Primers used for ChIP assay.

### Electron microscopy

In order to visualize the mitochondria, the kidney tissues of SHR and WKY were fixed with 3% buffered glutaraldehyde. Subsequently, tissues were processed as described previously^30^, before viewing under JEM-1400 plus transmission electron microscope (Jeol, Japan).

### Statistical analysis

All the experiments were carried out at least three times and the statistical significance for each experiment was determined through unpaired, two-tailed Students t-test using Prism 5 program (GraphPad software, USA).

## RESULTS

### Endogenous expression of mitochondrial transcription factors is elevated in SHR kidneys

To estimate the expression of mitochondrial transcription factors (viz. Tfam, Tfb1m and Tfb2m) in SHR and WKY kidney tissues we analyzed the mRNA and protein levels of these genes by qPCR and Western blot analysis. SHR kidneys displayed elevated mtTFs’ mRNA levels (Tfam: ∼3.2-fold, p<0.001; Tfb1m: ∼2.1-fold, p<0.01 and Tfb2m: ∼2.4-fold, p<0.01; Figure 1A-C). Consistently, protein levels of mtTFs were elevated in SHR kidney as compared to WKY. Additionally, protein levels of subunits of the respiratory chain were higher in SHR kidney (Figure 1D-E).

**Figure 1.**
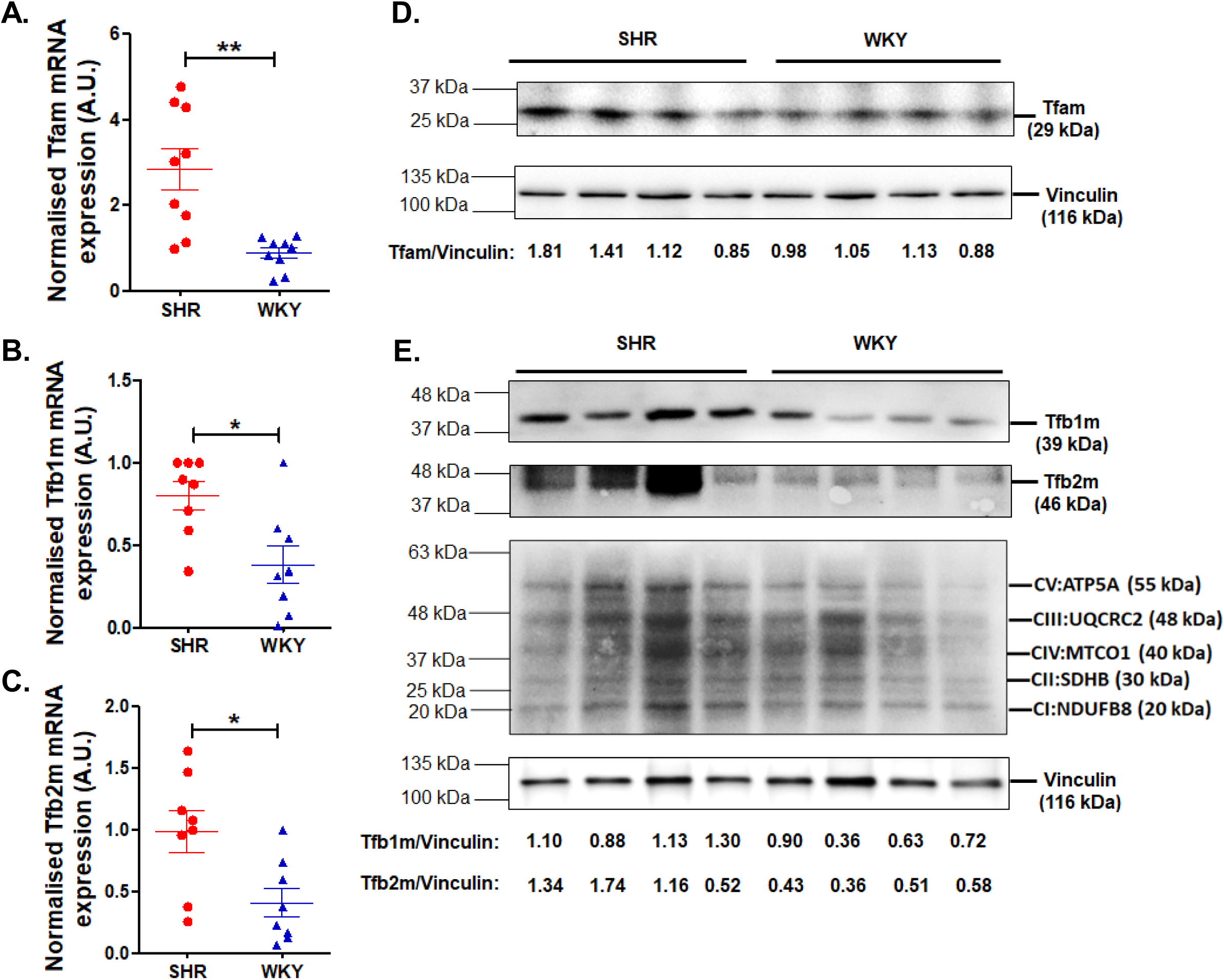
Endogenous expression of mitochondrial transcription factors in SHR/WKY kidney: Total RNA and protein were isolated from kidney tissue homogenates of SHR (n=10) and WKY (n=10) animals. RNA was subjected to reverse transcription and 40 ng of cDNA was used as a template for qPCR to estimate endogenous transcript levels of (A) Tfam, (B) Tfb1m and (C) Tfb2m using gene specific primers. Normalisation was carried out with respect to expression of β-Actin. Unpaired, 2-tailed Student’s t-test was used for statistical analysis. *p<0.05. Western blot analysis was performed using kidney homogenates from SHR and WKY. Representative images are shown for (D) Tfam, (E) Tfb1m, Tfb2m and OXPHOS. Vinculin was used for normalization of representative Western blot images and the densitometric quantifications are shown below the images.

### PGC-1α and HIF-1α expressions are elevated in SHR kidneys

Since the mtTFs expression was elevated in SHR, we determined the expression of PGC-1α, the master regulator of mitogenesis. Indeed, the expression of PGC-1α was ∼4-fold (p<0.0001) higher in SHR kidneys compared to WKY (Figure 2A). Furthermore, we observed correspondingly higher protein levels of PGC-1α in SHR as compared to WKY (Figure 2B).

**Figure 2.**
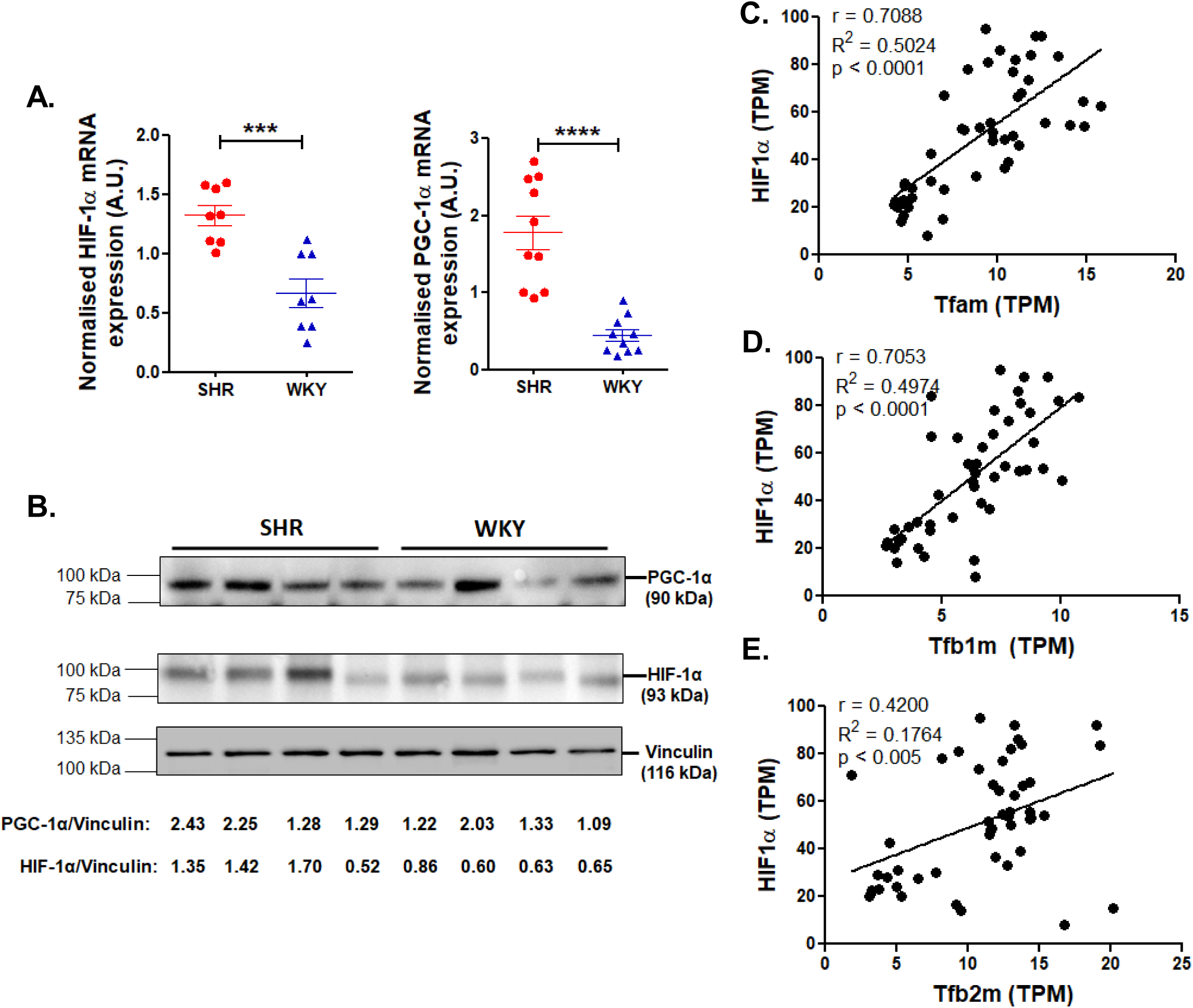
Endogenous expression of PGC-1α and HIF-1α in SHR/WKY kidney tissues and positive correlation between HIF-1α and mtTFs expression in human tissues: Total RNA and protein were isolated from kidney tissue homogenates of SHR (n=10) and WKY (n=10). (A) RNA was subjected to reverse transcription and 40 ng of cDNA was used in qPCR to estimate endogenous transcript levels of HIF-1α and PGC-1α using gene specific primers. The gene expression in each case was normalised to the expression of β-Actin expression. Unpaired, 2-tailed Student’s t-test was used for statistical analysis. ****p<0.0001 and ***p<0.001. (B) Western blot analysis was carried out using kidney homogenates from SHR and WKY. Representative images are shown for PGC-1α and HIF-1α protein levels. Vinculin was used for normalization of representative Western blot images and the densitometric quantifications are shown below the images. Correlation between the transcript levels of HIF-1α and (C) Tfam, (D) Tfb1m, (E) Tfb2m in various human tissues was plotted using transcriptomic data from the GTEx portal (n=48). TPM = transcripts per million.

To understand whether increased oxygen consumption rates in SHR^4, 31^ causes intracellular hypoxia, we estimated the expression of Hypoxia-Inducible-Factor-1-alpha (HIF-1α), an indicator of cellular oxygen deficiency. Interestingly, the mRNA expression of HIF-1α was higher in SHR kidneys by ∼2-fold (p<0.0002) as compared to WKY (Figure 2A). and the protein levels were also elevated (Figure 2B).

### Positive correlation between HIF-1α and mtTFs in various human tissue samples

In view of our finding that mtTFs and HIF-1α are concomitantly elevated in SHR (as compared to WKY) we analyzed the expression of these genes in various human tissues using RNAseq data from the GTEx portal^32^. Each of the mtTFs displayed significantly positive correlation with HIF-1α (viz. Tfam and HIF-1α, Pearson’s *r* = 0.7088, p < 0.0001; Tfb1m and HIF-1α, Pearson’s *r* = 0.7063, p < 0.0001 and Tfb2m and HIF-1α, Pearson’s *r* = 0.3962, p = 0.0048; Figure 2C-E).

### Effect of acute hypoxia on the promoter activity, mRNA and protein levels of mtTFs

Since hypoxic SHR kidneys display elevated levels of mtTFs (Figure. 1), we aimed to elucidate hypoxia-mediated expression of mtTFs in an *in vitro* cell culture model. Accordingly, we transfected NRK52e rat kidney epithelial cells with mtTFs’ promoter-luciferase reporter plasmids and subjected those to two hours of acute hypoxic stress 12 hours post-transfection. Hypoxia caused significant enhancements in the promoter activity of each mtTF (Tfam: ∼12-fold, p<0.01; Tfb1m: ∼3.6-fold, p<0.001; Tfb2m: 1.7-fold, p<0.05; Figure 3A, 3B, 3C). Furthermore, qPCR analysis showed elevated transcript levels of mtTFs (Tfam: 1.4-fold, p<0.05; Tfb1m: 1.5-fold, p<0.05 and Tfb2m: 2.8-fold, p<0.05; Figure 3D, 3E, 3F) under acute hypoxia. Similarly, acute hypoxia elevated the protein levels of mtTFs (Figure 3G). Successful induction of hypoxia was confirmed by elevated HIF-1α protein levels. Interestingly, OXPHOS subunits and PGC-1α levels were also higher during hypoxic conditions (Figure 3G).

**Figure 3.**
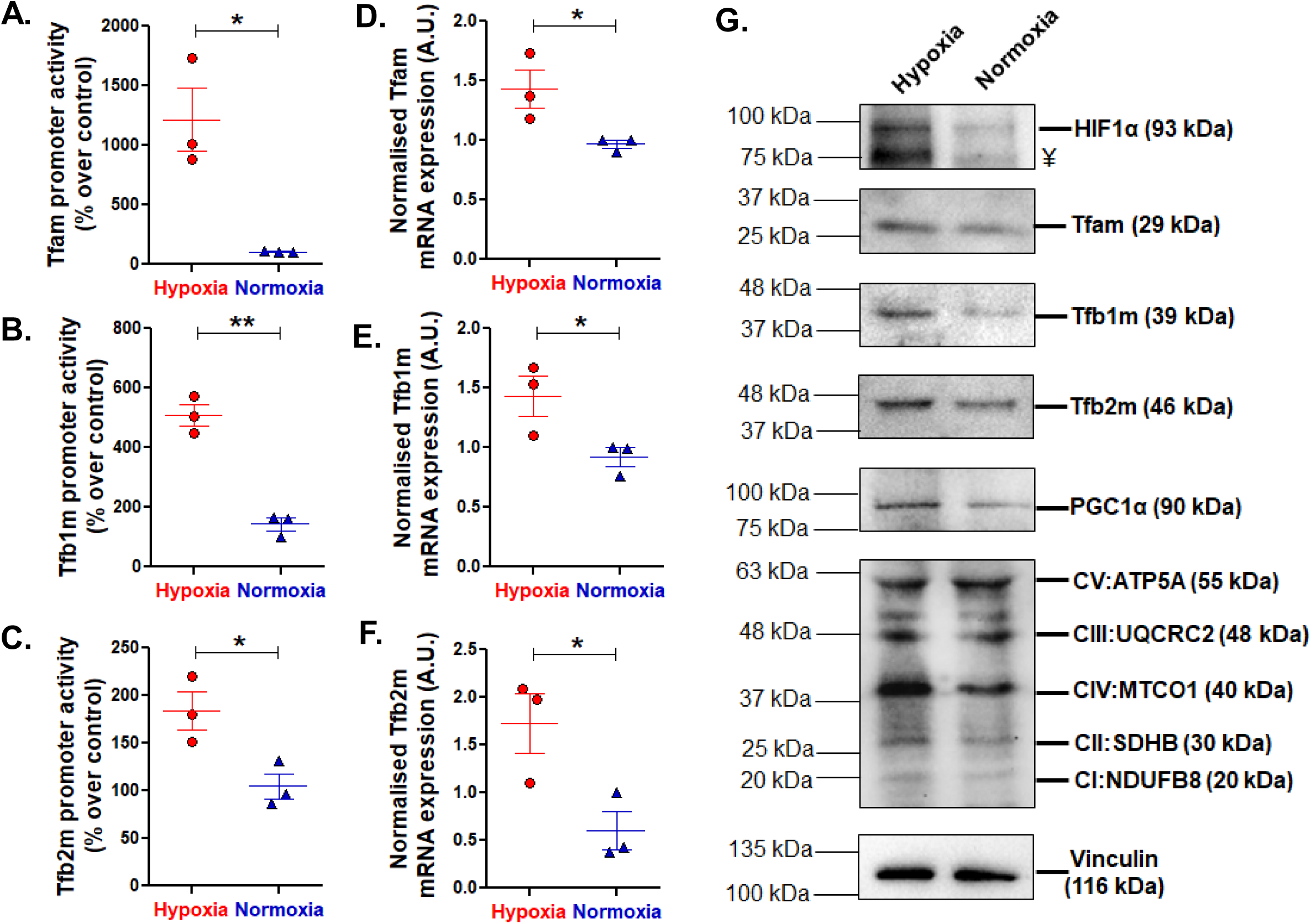
Effect of acute hypoxia on the promoter activity, mRNA and protein expression of mitochondrial transcription factors in cultured rat kidney epithelial cells: (A-C) NRK52e cells transfected with promoter luciferase-reporter constructs of (A) Tfam, (B) Tfb1m or (C) Tfb2m were subjected to acute hypoxia for two hours and luciferase activity was measured. Results are expressed as percentage over control after normalisation. mRNA expression of (D) Tfam, (E) Tfb1m or (F) Tfb2m were estimated through qPCR using RNA isolated from NRK52e cells subjected to acute hypoxia or normoxia. Expression of β-Actin was used for normalization. (G) Representative Western blot images showing protein levels of HIF-1α, Tfam, Tfb1m, Tfb2m, PGC-1α and total OXPHOS after acute hypoxia are shown. Vinculin was used as a normalization control. Student’s t-test (unpaired, two-tailed) was used for determining the statistical significance of the changes in expression after hypoxia. *p<0.05 and **p<0.001 with respect to the normoxic condition. ¥ indicates the position of the degradation product of HIF-1α seen at 75 kDa.

### Identification of multiple HIF-1α binding sites in the promoter region of mtTFs and their conservation across mammals

Acute hypoxia led to an increase in the expression of mtTFs at both transcriptional and translational levels. In addition, RNA-seq data from GTEx portal revealed a strong positive correlation between HIF-1α and mtTFs in human tissues. Therefore, we probed for the presence of HIF-1α binding sites (consensus matrix, Figure 4A) in the promoter region of mtTFs using *in silico* tools. Multiple such tools predicted several binding sites for HIF-1α within the proximal promoter of mtTFs (Table S1). Interestingly, at least one HRE (with good presiction scores, Figure 4A) from each promoter was conserved across various mammals suggesting a probable evolutionary importance of these HREs (Figure 4B-D).

**Figure 4.**
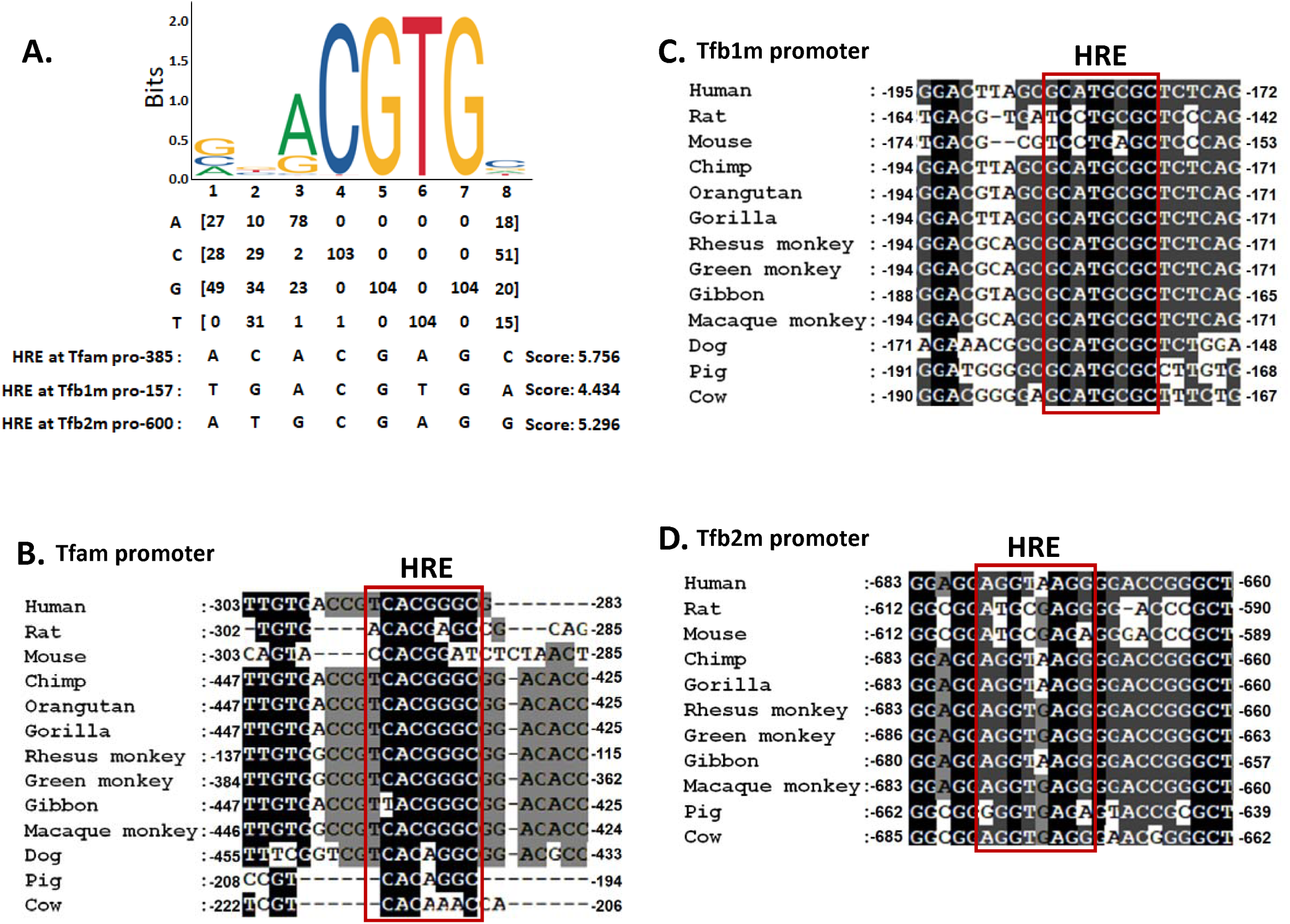
Conservation of HIF-1α binding sites in the promoter regions of mitochondrial transcription factors across mammals: (A) Position weight matrix for HIF-1α obtained from JASPAR database is shown. The promoters of mitochondrial transcription factors were analysed *in silico* using various online tools for the presence of Hypoxia Response Elements (HREs). The promoter sequences of rat Tfam, Tfb1m and Tfb2m harbouring the predicted HREs are shown below the matrix. The score for each of these sites is indicated for each promoter sequence. Conserved HREs predicted for these promoters across various mammals for (B) Tfam, (C) Tfb1m and (D) Tfb2m promoter sequences are shown. Red boxed region indicates the conserved HRE.

### *In vivo* interactions of HIF-1α with promoters of Tfam, Tfb1m and Tfb2m in the context of chromatin

We found four Hypoxia Response Elements (HREs) in Tfam, two each in Tfb1m and Tfb2m promoters, respectively (Figure 5A). In order to assess which of the predicted HIF-1α binding sites are functional, we carried out chromatin immunoprecipitation assays. qPCR was performed using immunoprecipitated DNA (ChIP grade HIF-1α antibody) from NRK52e cells cultured under normoxia/acute hypoxic conditions. Primers were synthesized to span one or more of these HREs in the promoter regions of mtTFs as shown in Figure 5A. Our analysis revealed that acute hypoxic stress enhanced binding of HIF-1α to HRE1/2 and HRE3 of Tfam promoter (∼3-fold, p<0.01 and ∼1.7-fold, p<0.001 respectively), HRE1 of Tfb1m promoter (∼2.6-fold, p<0.001), and HRE2 of Tfb2m promoter (∼2.5-fold, p<0.01). However, there was no significant difference in HIF-1α binding to HRE4 of Tfam, HRE2 of Tfb1m and HRE1 of Tfb2m (Figure 5B). Of note, our conservation analysis show that the HREs conserved across mammals are all functional.

**Figure 5.**
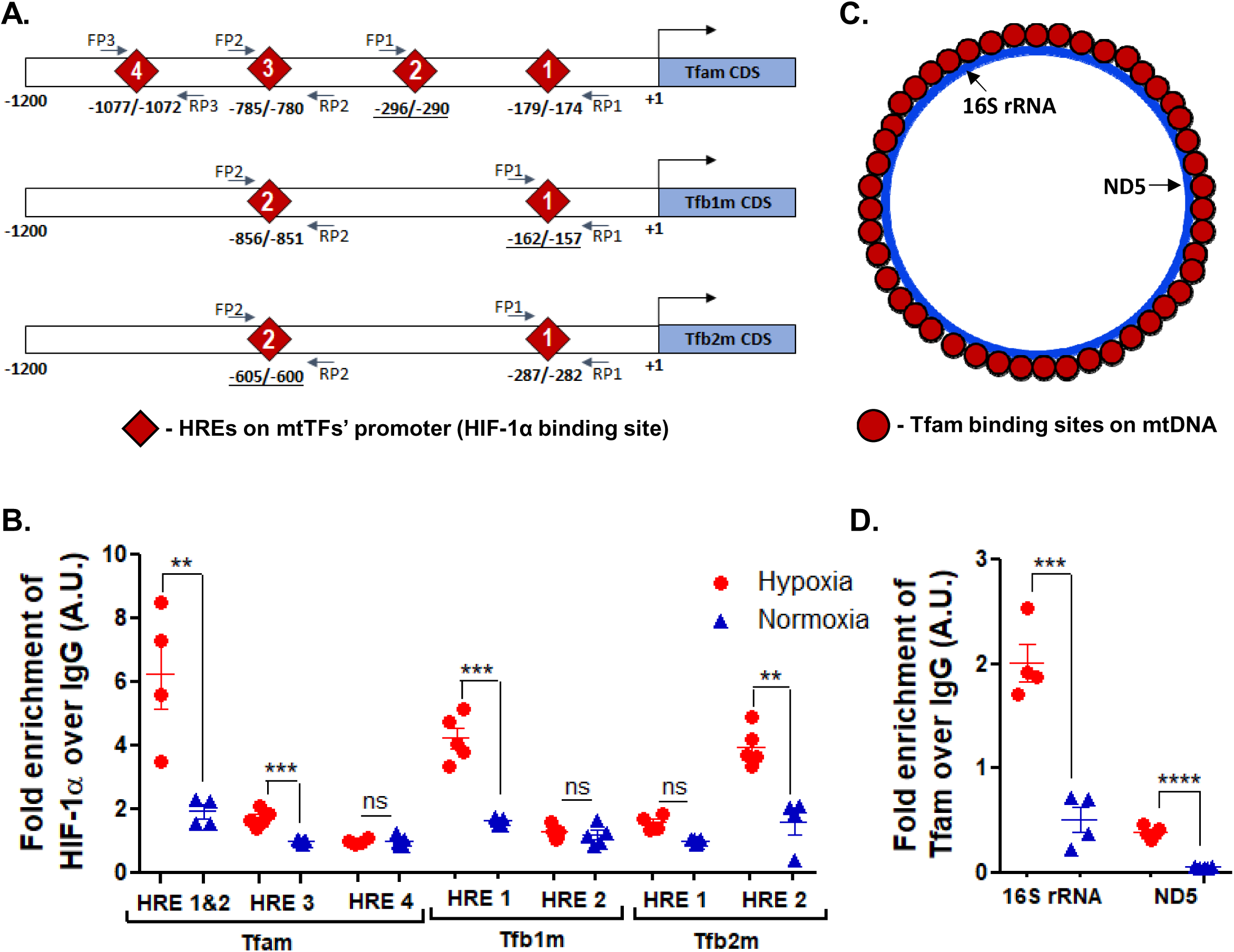
Interaction of HIF1α and Tfam with mtTFs promoters and mtDNA, respectively: (A) Schematic representation of HIF-1α binding sites on the promoters of mtTFs. The primers flanking each of these binding sites are indicated as arrows. Bold and underlined sites denote conserved HREs. (C) Schematic representation of Tfam binding sites on mtDNA. Positions of16S rRNA and ND5 genes are indicated. NRK52e cells were subjected to acute hypoxia for two hours. Chromatin was reverse crosslinked and fragmented prior to immunoprecipitation with antibodies specific to HIF-1α and Tfam. The immunoprecipitated DNA was subjected to qPCR analysis using primers flanking (B) various HREs or (D) mtDNA genes to assess the binding of HIF-1α and Tfam respectively. Students t-test was used to determine the statistical significance of differential binding. **p<0.01, ***p<0.001 and ****p<0.0001 with respect to the normoxic condition. ns: not significant.

### Interactions of Tfam with mtDNA in cultured kidney cells

One of the functions of Tfam is to coat the entire mtDNA in a histone-like manner (schematically represented in Figure 5C) so as to protect it from ROS-induced damage^33^. In order to investigate the functional significance of enhanced expression of Tfam after hypoxia, we assessed differential binding of Tfam to mtDNA using ChIP assay. The immunoprecipitated DNA (using ChIP grade Tfam antibody) from NRK52e cells cultured under normoxic/hypoxic condition was used in qPCR reactions in order to determine if there is differential binding of Tfam to various regions of mtDNA. Primers were designed flanking the 16S rRNA and ND5 genes to test for Tfam binding. We observed that Tfam binding to these regions is significantly enhanced (16S rRNA: ∼3-fold and ND5: ∼5-fold; p<0.001) upon exposure to acute hypoxia (Figure 5D).

### Functional consequence of increased expression of mitochondrial transcription factors

Since, HIF-1α directly mediates transcriptional activation of mtTFs during acute hypoxia (Figure 5), we explored the status of replication and transcription rates of mtDNA under these conditions. Accordingly, mtDNA and mtRNA were quantified in NRK52e cells subjected to acute hypoxia through qPCR analysis. We chose three genes: 16S rRNA, ND5 and ND6 transcribed from HSP1, HSP2 (Heavy Strand Promoter 1,2) and LSP (Light Strand Promoter) promoters of mtDNA, respectively^21^. Our analysis showed augmented mtDNA copies as evident from enhanced expression of 16S rRNA (∼2.2-fold, p<0.05; Figure 6A), ND5 (∼4-fold, p<0.001; Figure 6B) and ND6 (∼4-fold, p<0.0001; Figure 6C) after oxygen deprivation. Consistent with mtDNA amounts, we observed significant enhancements in mtRNA apparent from the expression of 16S rRNA (∼1.7-fold, p<0.01; Figure 6D), ND5 (∼3.3-fold, p<0.001; Figure 6E) and ND6 (∼3-fold, p=0.08; Figure 6F). Thus, acute hypoxia seems to augment the rates of replication and transcription of the mitochondrial genome, which in turn may enhance the protein translation within mitochondria (Figure 3G).

**Figure 6.**
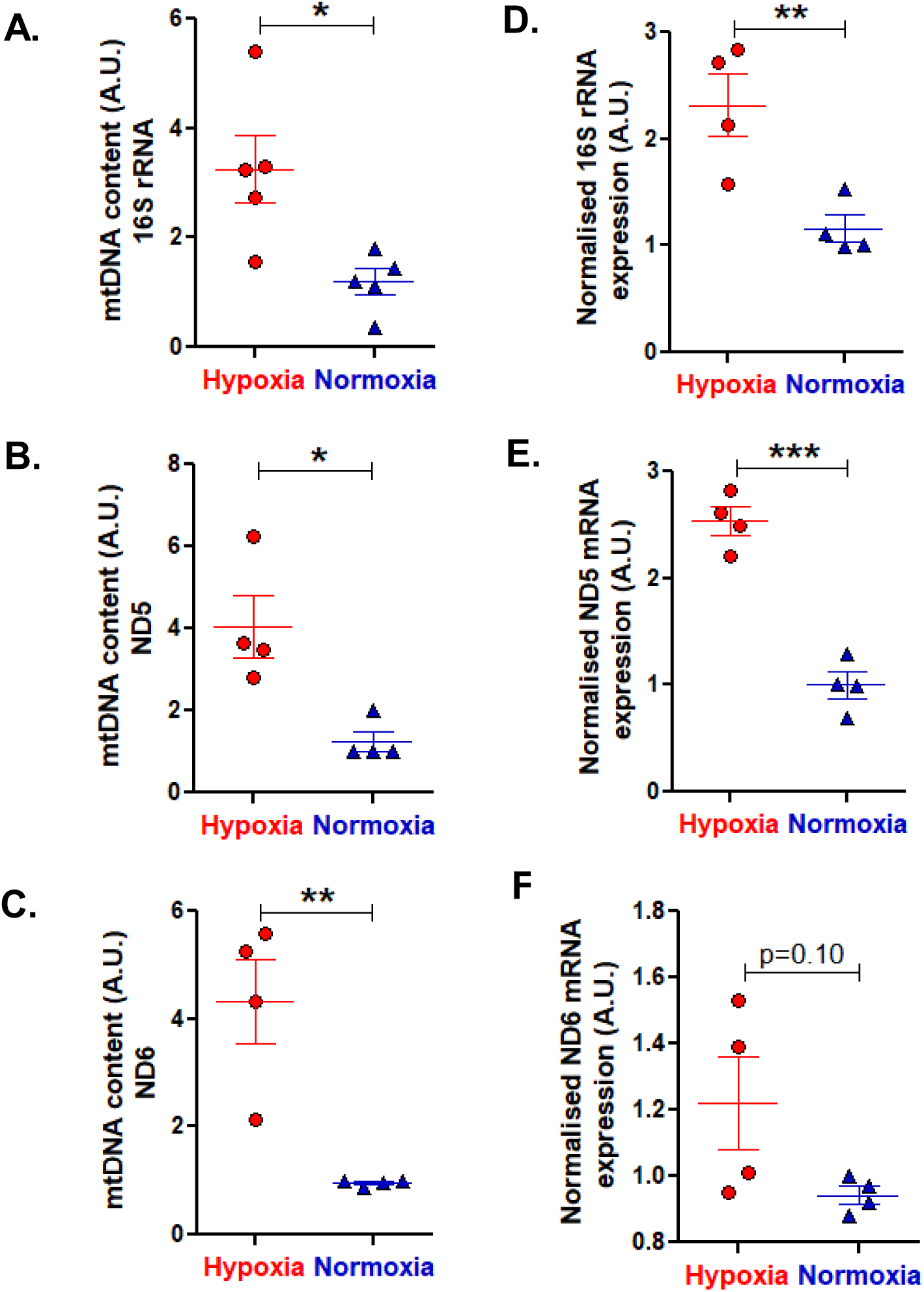
Effect of acute hypoxia on the expression of mtDNA and mtRNA in cultured kidney cells: NRK52e cells were subjected to acute hypoxia for two hours and DNA, RNA and protein were isolated. qPCR was performed using genomic DNA as template and primers specific for (A) 16S rRNA, (B) ND5 and (C) ND6 and normalised with 18S rRNA. Similar analysis was performed using cDNA as template and normalised with β-Actin to estimate mtRNA (D) 16S rRNA, (E) ND5 and (F) ND6. Student’s t-test (unpaired, two-tailed) was used for determining the statistical significance of the changes in expression after hypoxia. *p<0.05, ** p<0.01 and ***p<0.001 with respect to the normoxic condition.

### Hypertensive rats exhibit increased mitochondrial gene expression

In order to test whether acute hypoxia increases mtDNA and mtRNA in an animal model of genetic hypertension (in line with our *in vitro* finding; Figure 6), we isolated total DNA and total RNA from the kidney tissues of SHR and WKY for expression analysis by qPCR. mtDNA levels. mtDNA levels are elevated for 16S rRNA, ND5 and ND6 up to ∼3-fold in SHR (p<0.01, p=0.05 and p<0.05, respectively, Figure 7A, 7B, 7C). Furthermore, by performing qPCR analysis using total RNA, we demonstrate that the mtRNA are significantly higher in SHR kidney tissues in contrast to WKY (16S rRNA: ∼1.8-fold, p<0.01; ND5: ∼3.8-fold, p<0.05; ND6: ∼3.6-fold, p = 0.07; Figure 7D, 7E, 7F). Therefore, hypoxic environment in SHR kidneys influence the expression of their mitochondrial genome when compared to the kidneys of WKY rats.

**Figure 7.**
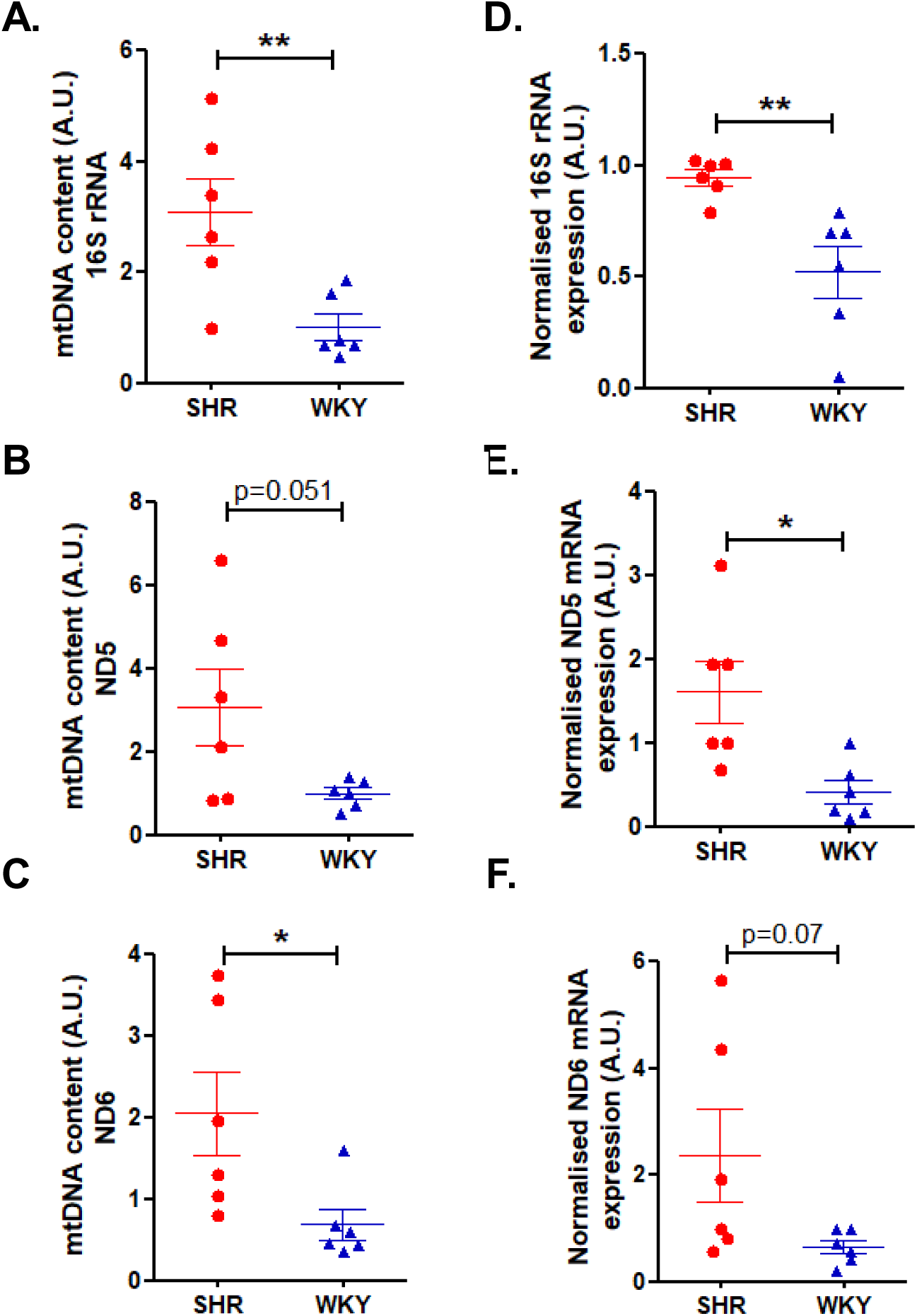
Expression of mtDNA and mtRNA in kidney tissues of SHR and WKY: qPCR was performed using primers specific for three mitochondrial genes using genomic DNA from SHR (n=6) and WKY (n=6) kidney tissue homogenates as template and primers specific for (A) 16S rRNA, (B) ND5 and (C) ND6. Similar analysis was performed using cDNA from SHR and WKY kidney tissue lysates as template and primers specific for (D) 16S rRNA, (E) ND5 and (F) ND6. mtDNA levels were normalized with respect to the abundance of 18S rRNA and mtRNA was normalized with respect to β-Actin expression. Student’s t-test (unpaired, two-tailed) was used to determine the statistical significance of differential expression. *p<0.05, **p<0.01 with respect to WKY.

### Mitochondrial abundance in SHR and WKY kidneys

Since SHR kidneys (that are known to be oxygen deprived) express elevated levels of mtTFs, mtDNA, mtRNA, PGC1-α and OXPHOS proteins as compared to the age-matched WKY control animals, we tested if there are differences in the mitochondrial abundance in the kidney tissue of these animals. Our TEM analysis revealed that the mitochondrial abundance seems to be similar between SHR and WKY kidney tissues (Figure 8).

**Figure 8:**
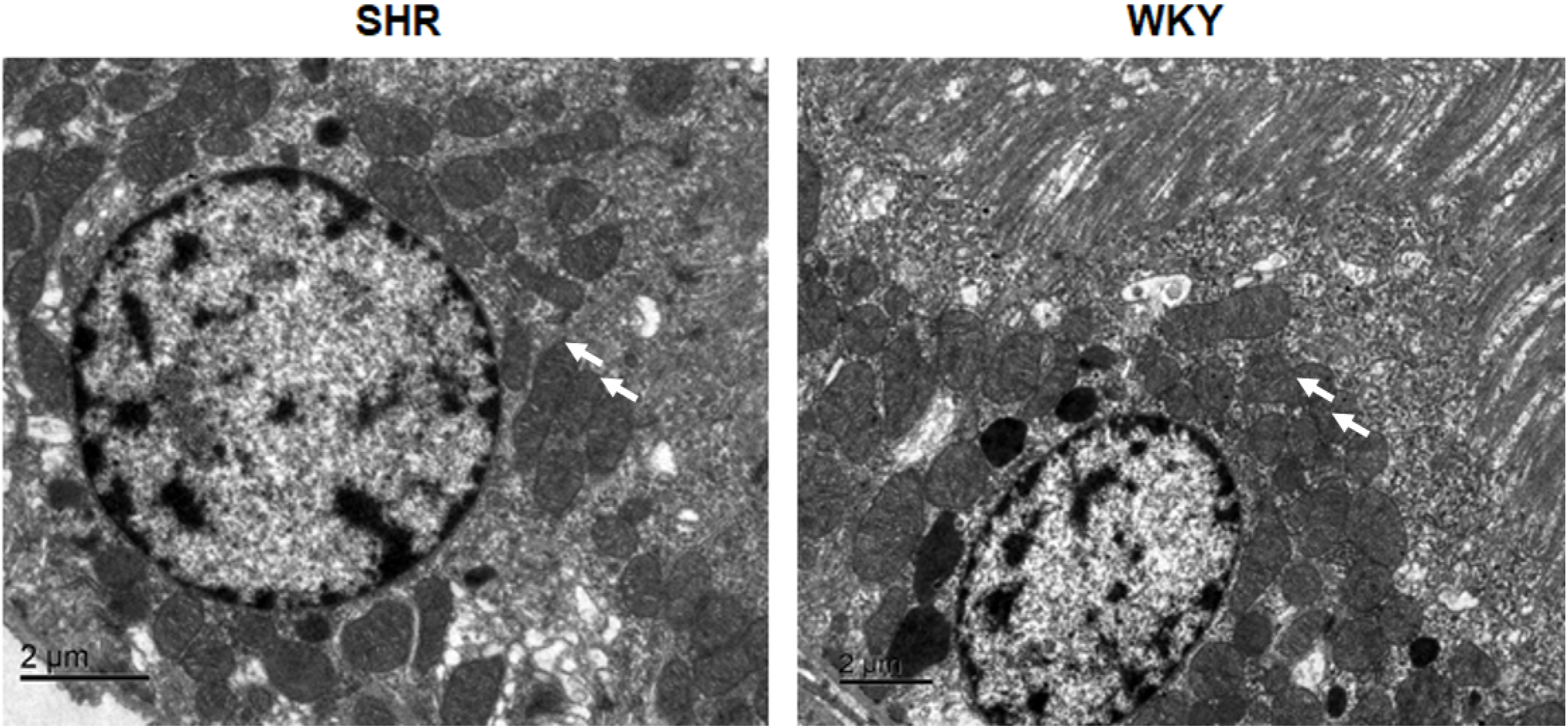
Mitochondrial ultrastructure in the renal cortex of SHR and WKY: Kidney tissues of SHR and WKY were fixed by vascular perfusion using 3% buffered glutaraldehyde. The renal cortices were excised into small pieces and post-fixed with 1 % osmium tetroxide. Subsequently, the tissues were dehydrated with increasing concentration of alcohol and cleared. Finally, infiltration and embedding were done using araldite. Ultrathin sections were cut using ultramicrotome (UC7, Leica) and stained for viewing under TEM. Numerous mitochondria (white arrows) were seen in tubular epithelial cells of both SHR and WKY kidney (Scale bar 2 µm).

## DISCUSSION

### Overview

Tubulo-interstitial Hypoxia (TiH) is a major contributing factor for many kidney diseases viz. Chronic Kidney Disease (CKD), Acute Kidney Injury (AKI), End Stage Renal Disease (ESRD), etc. Interestingly, TiH has been implicated in hypertension-induced renal malfunction and other types of nephropathy^4^. Of note, TiH is often accompanied by mitochondrial dysfunction^34, 35^. Particularly, TiH is anteceded by oxygen deficiency resulting from increased oxygen consumption by mitochondria of renal tubular epithelial cells^4^. However, the effect of renal TiH on mitochondrial biogenesis is partially understood. In this study, we used a cell culture model of acute hypoxia and a rodent model of hypertension (SHR and WKY) to gain insights about mitogenesis during TiH. Our findings (summarized in Figure 9) indicate that the expression of crucial mitochondrial proteins encoded by the nuclear and mitochondrial genome is higher in the hypoxic kidneys of SHR in contrast to WKY and also during acute hypoxia *in vitro* as compared to normoxic conditions. Interestingly, the kidneys of SHR and WKY are similar in their mitochondrial abundance (Figure 8).

**Figure 9:**
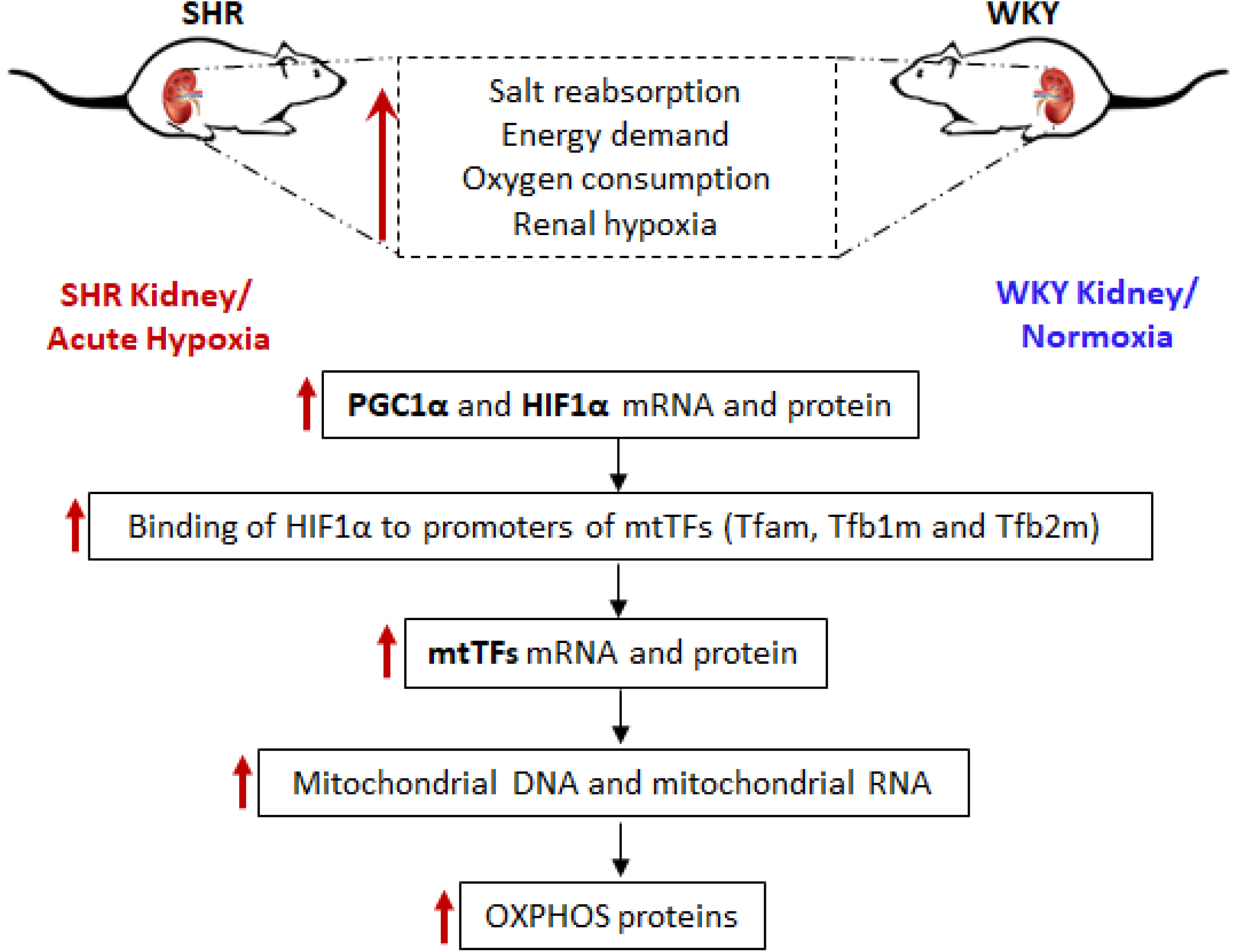
Plausible mechanisms of regulation of mitochondrial transcription factors during acute hypoxia and in hypertensive renal physiology. Hypoxic SHR kidneys and NRK52e cells subjected to acute hypoxia display augmented expression of mitochondrial transcription factors. HIF-1α mediates the increase in expression of mtTFs at the transcriptional level. Augmented mtTFs result in increased mtDNA and mtRNA levels leading to enhanced OXPHOS expression. Increased OXPHOS may improve mitochondrial function and maintenance in hypoxic kidneys of young SHR when compared to relatively normoxic WKY kidneys.

### Kidneys of young, pre-hypertensive SHRs exhibit augmented mitochondrial proteins

Kidneys are second to the heart in energy consumption and exhibit a high resting metabolic rate, and thereby, requiring excessive energy for efficient functioning^36, 37^. Moreover, renal tubular epithelial cells have low glycolytic capacity and rely majorly on fatty acid oxidation (FAO) as source of energy^38^. Thus, proper mitochondrial function is essential in order to maintain adequate salt-water reabsorption. TiH has been observed in the hypertensive rodents (SHR) as opposed to the normotensive ones (WKY). Increased TiH has been proposed to arise from decreased oxygen utilization efficiency in SHR^39^. Additionally, SHR kidneys have been shown to carry out increased sodium reabsorption^40^.

In this study, we initially focused on mitochondrial biogenesis in SHR and WKY since mitochondrial health is imperative for active solute transport within the nephrons. To this end, we analyzed the expression of genes indicative of mitochondrial activity and abundance (viz. PGC-1α, mtTFs) in the kidneys of SHR and WKY (Figure 1, 2A and 2B). We observed that the renal abundance of PGC-1α and mtTFs were significantly higher in SHR as compared to WKY. Correspondingly, the expression of mtDNA, mtRNA and electron transport chain (ETC) subunits were also elevated in the SHR kidneys in comparison to WKY (Figure 7). Interestingly, our TEM analysis revealed that mitochondrial abundance was not significantly different in the renal cortex between the two rat strains (Figure 8). Several studies provide physiological evidence of greater ATP demand in SHR kidneys. For example, it has been reported that the activity of Na^+^/K^+^-ATPase pumps in renal cells of pre-hypertensive rats is higher than the normotensive counterparts^41^. Moreover, perfusion of isolated kidneys from 3-month-old SHR and WKY animals revealed higher sodium reabsorption in SHR kidney^40^. Multiple studies report pathological increases in salt and water reabsorption by SHR kidneys^42–45^. In addition, efficiency of sodium transport (total sodium transported per molecule of oxygen consumed) is compromised in SHR, adding to ATP demand^14^. Elevated mitochondrial proteins could thus be purported to aid in ATP synthesis.

Our observations are in line with another study that demonstrated increased mitochondrial activity (as shown by augmented expression and activity of pyruvate dehydrogenase, enhanced flux of acetyl-CoA and increased ATP synthesis) in young SHR compared to WKY^13^. Our results and findings from these studies suggest that renal epithelial cells from young, pre-hypertensive SHR exhibit enhanced mitochondrial function as compared to the normotensive WKY rats.

### Positive correlation between HIF-1α and mtTFs transcripts in multiple tissues

We observed that HIF-1α expression in the renal tissues of SHR was higher when compared to WKY (Figure 2A, 2B), indicating local oxygen deprivation, probably arising from increased oxygen consumption in the tissues of SHR^14^. This is consistent with another study reporting that HIF-1α^46^ is higher in the renal medulla where the tubular cells have to work against a greater concentration gradient in order to transport sodium^47^. The increased oxygen consumption in SHR kidneys has also been attributed to inefficiency in oxygen utilization and nitric oxide bioavailability^39^. Our observation that HIF-1α expression is elevated in SHR as compared to WKY, suggests greater oxygen deficiency in SHR. It has been reported that overexpression of HIF-1α ameliorates kidney oxygenation and improves mitochondrial function in a rodent model of subtotal nephrectomy^34^. Therefore, the elevated expression of HIF-1α in SHR as compared to WKY may compensate for in-efficient oxygen utilization in these rats. Additionally, it has been reported that SHR kidneys have compromised antioxidant defenses^48^. Subsequent activation of cellular stress sensor Nrf2 (nuclear factor erythroid-2 related factor-2) might be responsible for transcriptional activation of HIF-1α in the kidneys of SHR animals^49^. Indeed, the activation of Nrf2 pathway has been proposed to prevent tubular damage to kidneys^50, 51^.

RNA-seq data for various human tissues from healthy individuals displays positive correlation between HIF-1α and mtTFs (Figure 2C-E) suggesting that HIF-1α might regulate the expression of mtTFs under normal physiological conditions. It is well established that high levels of AMP/ATP activate AMPK (AMP-kinase) which, in turn, triggers HIF-1α transcription^52^. Notably, AMPK activation is also indicative of energy demand of the cells. Therefore, it is possible that HIF-1α transcription is regulated based on the amount of oxygen consumed and that HIF-1α protein is stabilized under oxygen-deficient conditions. A positive correlation is, thus, understandable since mitochondrial transcription factors are directly related to ATP production and oxygen consumption. Moreover, it has been reported that under physiologically normal oxygen concentrations, PGC-1α overexpression leads to intracellular oxygen deficiency and stabilizes HIF-1α in skeletal muscles. HIF-1α further activates its downstream targets and prepares the cells for oxygen deficiency arising from enhanced oxygen consumption^2^.

### Acute hypoxia enhances the expression of mitochondrial transcription factors via HIF-1α

Data from rodent models of hypertension indicated a hypoxic environment in the SHR kidney. We also observed a strong positive correlation between HIF-1α and mtTFs. Next, we asked whether enhanced HIF-1α levels modulate the expression of mtTFs. In this direction, we sought to study the transcriptional regulation of mtTFs under conditions of acute hypoxia. Indeed, the mtTFs’ promoter activities were increased upon hypoxic stress *in vitro.* A concomitant increase in the mRNA and protein expression of these genes was also observed (Figure 3). Upon further analysis, we observed elevated expression of mitochondrial DNA, RNA and OXPHOS proteins after acute hypoxia (Figure 6). These results indicate increased mitochondrial replication, transcription and translation during acute hypoxia. Additionally, we found enhanced expression of PGC-1a upon acute hypoxic stress.

### HIF-1α interacts with promoters of mtTFs to enhance their gene expression

In line with our *in-silico* predictions, ChIP analysis showed that HIF-1α directly binds to the promoters of mtTFs on at least one of the predicted sites during hypoxia (Figure 5A, 5B). Furthermore, these binding sites are conserved across mammals (Figure 4). Hence, it is possible that direct induction of mtTFs expression by HIF-1α might occur in other mammals, suggesting that it is an evolutionarily crucial regulatory phenomenon. Therefore, our results suggest that HIF-1α-mediated enhanced expression of mtTFs during acute hypoxia might promote expression of mitochondrial genome in order to increase energy production.

### Tfam binds to mtDNA during acute hypoxia

Mitochondria are major sites of ROS production and it is well established that hypoxic stress increases ROS levels within these organelles. ROS are known to cause DNA damage of various types. Studies have established that mtDNA is found in the form of nucleoids. Tfam is a major constituent of these nucleoids and is said to protect mtDNA from ROS-induced damage^5, 50^, apart from regulating the replication and transcription rates. Our ChIP assays reveal that hypoxia enhances the binding of Tfam to multiple sites on mtDNA (Figure 5C-D), probably to prevent oxidative damage to mtDNA during acute hypoxia. Such observations have been made previously^53^. However, if the DNA is compacted, how would replication and transcription occur? A single mitochondrion is known to contain many copies of its genome. Tfam binding to mtDNA is cooperative in nature, i.e., Tfam binds with greater affinity to DNA that already is bound to other molecules of Tfam^54^. We speculate that out of multiple copies of the genome, most of it might be compacted while some still serve to enable gene expression. This kind of regulation is advantageous since if uncompacted mtDNA is damaged, there are still many more mtDNA copies to take its place enabling the cells to delay the event of severe mitochondrial dysfunction. A similar observation has been made, where it was shown that oxidative modifications to mtDNA (due to acute hypoxia) promotes Tfam binding to mtDNA and enhances its replication in pulmonary arterial endothelial cells has been reported^55^. This is in line with our observation that mtDNA copies increase after acute hypoxia. We also observe concomitant increases in mitochondria-encoded RNA.

### Conclusions and perspectives

This study shows, for the first time to our knowledge, that acute hypoxia enhances the transcription of mitochondrial transcription factors by direct interaction of HIF-1α with the promoters of these genes. The increase in HIF-1α levels after acute hypoxia is accompanied by an increase in PGC-1a level implicating that under our experimental conditions, mitochondrial biogenesis might be enhanced. Interestingly, Tfam binding to mtDNA is enhanced upon hypoxic stress. Additionally, our data indicates that short-term hypoxia increases the mitochondrial genome expression. These data are summarized in Figure 9.

In line with the *in vitro* data, the hypoxic SHR kidney (as revealed by enhanced expression of HIF-1α) exhibits higher expression of mtTFs and PGC-1α than WKY. Concomitant increases in mtDNA and mtRNA are also observed in SHR. This indicates heightened ATP requirements and increased oxygen consumption in SHR as compared to WKY. It should be noted that the animals used in our study are young, aged 4-6 weeks when SHRs are still pre-hypertensive. We propose that during early stages of hypertension, the organism’s physiology still functions to meet the increased energy demand and tries to keep the kidneys functional up to the best possible levels. Nevertheless, at a later age, ROS accumulation promotes inflammatory conditions that might be responsible for reduction in mitochondrial proteins observed in renal tissue of SHR as compared to WKY^56, 57^.

## ACKNOWLEDGEMENTS

This work was supported by a grant from the Science and Engineering Research Board (SERB), Department of Science and Technology, Government of India to NRM (project number EMR/2017/004250). Various Government of India Research fellowships were received from Council of Scientific and Industrial Research (BN), Department of Science and Technology (VA), Ministry of Human Resource Development (AAK) and Indian Council of Medical Research (SSR).

