## Supplemental data for "Hypoxia-mediated regulation of mitochondrial transcription factors: Implications for hypertensive renal physiology"

Running title: Regulation of mitochondrial transcription factors under acute hypoxia conditions

**Abbreviations**

mtTFs, mitochondrial transcription factors; OXPHOS, oxidative phosphorylation; ATP, adenosine triphosphate; ETC, electron transport chain, TiH, tubulointerstitial hypoxia; PGC-1alpha – peroxisome proliferator activated receptor gamma coactivator-1-alpha; SHR, spontaneously hypertensive rat; WKY, Wistar Kyoto rat; HIF-1α, hypoxia inducible factor-1α; HREs, hypoxia response elements; mtDNA, mitochondrial DNA; ChIP, chromatin immunoprecipitation.

**SUPPLEMENTARY METHODS**

**Rat strains**

In order to study the early changes in the hypertensive kidneys, the renal tissues from 4-6-week-old Spontaneously Hypertensive rats (SHR) and Wistar Kyoto normotensive rats (WKY) were harvested and stored in RNA-later solution to prevent RNA and protein degradation. For TEM analysis, the rats were perfused with 3% glutaraldehyde before dissection of the kidneys. The animal experiments were approved by the Institute Animal Ethics Committee at Indian Institute of Technology Madras.

**Generation of rat Tfam, Tfb1m and Tfb2m promoter-luciferase reporter constructs**

Promoter regions (~1.3 kb) of mitochondrial transcription factors were PCR-amplified using genomic DNA isolated from Wistar-Kyoto rats as template, primers specific to the promoters of Tfam, Tfb1m & Tfb2m (Table 1) and Phusion® High Fidelity DNA polymerase (Finnzymes, USA) at an annealing temperature of 62˚C. DMSO at a concentration of 3% was added to the reaction mixture. The purified PCR products and the pGL3Basic vector (Promega, USA) with firefly luciferase reporter were double digested with KpnI and XhoI restriction enzymes (New England Biolabs, UK). Ensuing this, the digested vector was incubated with Calf Intestinal phosphatase (CIP) (New England Biolabs, UK) for 15 minutes at buffer conditions specified by the manufacturer in order to remove 5’-PO_4_ overhangs. After additional purification of the inserts and vector, they were ligated using T4 DNA Ligase (ThermoFisher, USA). The ligation reaction was set at 10˚C in an insulated water bath for 12 hours so that the gradual increase in temperature during this time will let the ligase find an ambient condition to ligate the insert and vector. Ligation mixture was subsequently transformed into DH5α ultracompetent cells and the resulting colonies were screened to obtain the desired clones. Additionally, the clones were sequenced for confirmation.

***In silico* prediction of transcription factor binding sites in the mitochondrial transcription factors’ promoters**

This work aims to identify the hypoxia induced expression of mitochondrial transcription factors. Therefore, online transcription factor binding site prediction tools were used to specifically find binding sites for HIF1α, known as Hypoxia Response Elements (HREs), in the promoters of Tfam, Tfb1m and Tfb2m. The promoter sequences obtained from UCSC genome browser were given input sequences and HIF1α was specifically selected wherever possible. The scores predicted by the different tools for various HIF1α binding sites are listed in Table S1. The conservation of HREs across various mammalian species was analyzed. In order to accomplish this, the sequence of the mitochondrial transcription factors’ promoters from Human, Rat, Mouse, Chimp, Orangutan, Gorilla, Rhesus Monkey, Green Monkey, Gibbon, Macaque, Dog, Pig and Cow were obtained from UCSC genome browser or Ensembl. The GenBank/Ensembl accession IDs for these species are given in Table S2. These sequences were aligned using ClustalW and the alignment file was analyzed using the GeneDoc software.

**Cell culture and transfection**

Authenticated Normal Rat Kidney epithelial cells (NRK 52e) were obtained from the Indian national cell-line and hybridoma repository at the National Center for Cell Sciences, Pune, India. Dulbecco’s modified Eagle’s medium with high glucose and glutamine (HyClone, USA) supplemented with 10% fetal bovine serum, streptomycin (100 mg/mL) and penicillin G (100 U/ml) (Invitrogen, USA) was used for culturing cells in 25-cm^2^ tissue culture flasks, 60 mm or 100 mm dishes as described previously^1^. The cells were periodically tested for mycoplasma contamination through a PCR-based method. If contaminated, the cells were treated for three weeks with BM cyclins (Roche, USA) to get rid of contamination prior to performing our *in vitro* experiments. Transfections were done transiently in a 24-well plate using a peptide-based transfection reagent, Targefect F2 (Targeting Systems, USA), at a ratio of 1:1 wt/vol (DNA: Targefect F2). In each experiment, at ~70-80% confluency, 500 ng/well of Tfam-pro, Tfb1m-pro or Tfb2m-pro promoter-reporter construct was co-transfected with 250 ng/well of β-galactosidase expression plasmid serving as an internal control for transfection efficiency. Subsequently, the cells were subjected to acute hypoxia for two hours and lysed.

**Acute hypoxic exposure and luciferase assay**

12 h after transfection, cells were subjected to two hours of acute hypoxia. This was done by first placing the cells in a hypoxia chamber (BelArt, USA). Argon gas with 99% purity was infused for three mins to replace the existing oxygen to Argon. After this, the cabinet was sealed tightly and placed inside an incubator set to 37˚C. Parallelly, the cells to be kept under normoxic conditions were placed in the regular CO_2_ incubator. The incubation under hypoxia was for two hours, ensuing which they were lysed for downstream assays. Luciferase assays were carried out using a luciferase assay buffer (100 mM Tris-acetate, pH 7.8, 10 mM magnesium acetate, 1 mM EDTA, pH 8.0, 3 mM ATP and 100 µM beetle luciferin potassium salt) as described previously^1^. The values obtained as relative luciferase units per second (RLU/s) were normalized with β-galactosidase activity and total protein estimated by Bradford assay. Such experiments were repeated thrice, and the data are expressed as percentage over control of a representative experiment.

**RNA isolation and quantitative real-time PCR in NRK52e cells and rat model of essential hypertension**

In order to determine the expression profile of mitochondrial genes in our animal model, kidney tissues from SHR and WKY were isolated from young animals aged 4-6 weeks and stored in RNA later. These tissues or NRK52e cells (cultured under normoxic or hypoxic conditions) were washed with 1X PBS and homogenized using TRIzol reagent (Takara, USA). RNA was isolated using NucleoSpin RNA isolation kit (Takara, USA). The total RNA thus obtained was subjected to DNase treatment to remove any remnant DNA. Subsequently, cDNA was synthesised using cDNA synthesis kit (Takara, USA) according to manufacturer’s instructions. 20-40 ng of cDNA was used in each qPCR reaction and the fold change of expression with respect to the control (WKY normotensive rats or cells grown under normoxic condition) was calculated using the ΔΔCt method as described previously^1^. Primers specific for β-Actin was used as a house keeping control. The primers used for the mitochondrial genes profiled in this study are listed in Table 2.

**DNA isolation and estimation of mitochondrial DNA content**

In order to estimate mitochondrial DNA levels, the samples homogenised with Trizol for RNA isolation were used. After aspirating the aqueous phase containing RNA, 100% ethanol was added to the interphase and phenol-chloroform layer to precipitate DNA. Ensuing this, the samples were centrifuged at 200 rpm for 5 minutes at 4°C. The supernatant was discarded, and the pellet was washed twice with 0.1 M sodium citrate. The pellet obtained was then solubilised in DNase- and RNase- free water. Total DNA was then subjected to RNAse treatment to remove traces of RNA contamination. Total DNA containing both nuclear and mitochondrial DNA and was used in qPCR reactions to quantify amount of mtDNA as done previously^2^. The difference in the expression between SHR and WKY or normoxic vs hypoxic condition was calculated using ΔΔCt method using 18S rRNA as control.

**Western blotting for various proteins in NRK52e cells and rat model of essential hypertension**

Kidney tissues from SHR and WKY were isolated and washed with 1X PBS. The tissues were minced with RIPA buffer using a homogenizer. Sonication was performed at an amplitude of 30/s for 5 min in ice so as to breakdown the nucleic acids that might interfere with SDS-page and Western blotting. NRK52e cells were seeded in six well plate and exposed to hypoxia or normoxia for two hours. Supernatant was discarded and the cells were washed with 1X PBS prior to lysis using RIPA buffer. Lysates were subjected to gentle sonication at 30/s amplitude for 30 seconds. The proteins were then estimated using Bradford reagent (Bio-Rad, USA). 50 $\mu$g of protein was separated in a 10% SDS-polyacrylamide gel by electrophoresis. The separated proteins were transferred on to a PVDF membrane (Pall Life sciences, Mexico). Pre-stained protein ladder of wide range (Abcam, USA) was used as a molecular weight marker. The membranes were then blocked and probed with different concentrations of primary antibody: Tfam (CAT#sc28200; Santa Cruz Biotechnology, USA), Tfb1m (CAT#ARP34544_T100; Aviva, USA), Tfb2m (CAT#ab13676; Abcam, USA), OXPHOS cocktail (CAT#ab110413; Abcam, USA), HIF1α (CAT#ab463; Abcam, USA), PGC1α (CAT#ab54481; Abcam, USA) and Vinculin (CAT#V9131; Sigma-Aldrich, St. Louis, USA). After washing 3 times with 0.05% 1X TBST for 10 min, incubation was done with HRP conjugated secondary antibody raised against goat (ab6741; Abcam, USA), mouse (CAT#115-035-003; Jackson Immunoresearch, USA) or rabbit (CAT#111-035-003; Jackson Immunoresearch, USA). Washing steps were performed again 3 times with 0.05% 1X TBST for 5 min. The blocking agent, primary and secondary antibody dilutions and duration of incubation in each case is detailed in Table S3. The resulting chemiluminescence was detected using ECL substrate kit (Bio-Rad, USA). The band intensities were then quantified using ImageJ software.

**Chromatin immunoprecipitation**

ChIP assay was carried out as detailed earlier. Briefly, NRK52e cells were cultured in a 100 mm dish upto 90% confluency after which they were subjected to either hypoxia for two hours or not. Cells under both hypoxic and normoxic conditions were quickly washed with 1X PBS and crosslinked with 1% formaldehyde. Fixation was stopped using 0.125M glycine after which the cells were scraped and centrifuged at 3000 rpm for 7 min at 4˚C to pellet the nuclei. Nuclear pellet was then lysed with nuclei lysis buffer. Sonication was carried out at an amplitude of 35/s for 30 min in order to shear the chromatin. Sheared chromatin was precleared using beads that were blocked with BSA and E. coli genomic DNA. The precleared chromatin was incubated with 5 µg of HIF1α (CAT#ab463; Abcam, USA) or Tfam (CAT#sc28200; Santa Cruz Biotechnology, USA) antibody (in 1X ChIP buffer supplemented with protease inhibitors) overnight at 4˚C for immunoprecipitation. Parallelly, the chromatin was also incubated with mouse IgG (CAT# I5831; Sigma-Aldrich, St. Louis, USA) and rabbit IgG (CAT# I5006; Sigma-Aldrich, St. Louis, USA) antibody as a negative control to account for non-specific interactions during immunoprecipitation. Pull down of antibody bound DNA-protein complexes was done using ChIP grade G-sepharose beads (Invitrogen, USA). After stringent washing steps using low salt buffer, high salt buffer, lithium chloride and TE buffer as described previously, DNA was eluted and reverse crosslinked using 5M NaCl. Subsequently purified DNA was used for PCR amplification using primers that flank potential binding sites for HIF1α and Tfam. The list of primer sequences is given in Table 3.

**Electron microscopy**

In order to visualize the mitochondria in the renal tissue of SHR and WKY, 4-6-week-old rats were sacrificed and perfused with 3% glutaraldehyde in 0.1M phosphate buffer to fix the tissues. The kidneys were excised, and thin slices of the cortex and medulla were dissected and grossed into 1mm x 1mm bits and stored for an additional 24 hours in 3% glutaraldehyde. Tissue processing was done as described previously^3^. Briefly, following fixation the tissues were washed 3x15 minutes with 0.1M phosphate buffer. Post fixation was done using 1% Osmium tetroxide for 1.5 hours to enable fixation of lipids. The tissues were again washed 3x15 minutes with 0.1M phosphate buffer and dehydrated in increasing concentrations of distilled alcohol at 4°C. 70%, 80% and 90% ethanol were added successively for 1 hour each. Subsequently, to remove excess alcohol, propylene oxide was used as a clearing agent. Clearing was done 2x for 15 min. Next, the tissues were infiltrated with embedding medium by incubating them in a solution of propylene oxide: araldite Cy212 (1:1) overnight on a rotator at room temperature. Following this, the tissues were incubated for an additional 1 hour in a solution of propylene oxide: araldite Cy212 (1:2) and 5 hours in complete embedding medium. After successful infiltration, the tissues are placed on embedding molds containing araldite Cy212 and placed in an oven at 60°C to enable polymerization. Ultrathin sections were cut from tissue blocks using ultramicrotome (Leica EM UC6) and collected on copper grids. Ultrathin sections were contrasted with uranyl acetate and lead citrate and visualized under JEM-1400 plus transmission electron microscope (Jeol, Japan).

**SUPPLEMENTARY RESULTS**

**Table S1: HIF-1α binding sites predicted in the promoters of mitochondrial transcription factors using various *in silico* tools**

| **Gene** | **Location** | **Score** | | | | |
| --- | --- | --- | --- | --- | --- | --- |
|  |  | **Matinspector** | **ConSite** | **p-Match** | **Jaspar** | **LASAGNA** |
| **Tfam** | -174 | 0.75 | 5.75 | 0.80 | 15.61 | - |
|  | -385 | - | 7.10 | 0.80 | - | 9.73 |
|  | -780 | - | 5.75 | 0.80 | 6.18 | 2.80 |
|  | -1072 | - | 7.68 | - | 8.29 | - |
| **Tfb1m** | -157 | 1.00 | 4.43 | 0.80 | 4.40 | - |
|  | -851 | 0.75 | 4.89 | - | - | - |
| **Tfb2m** | -282 | 0.75 | 5.24 | 0.80 | - | - |
|  | -600 | 0.80 | 5.30 | 0.80 | - | - |

- Indicates data not available

**Table S2: GenBank/Ensembl accession IDs for mitochondrial transcription factors of mammals**

| **S. No** | **Species** | **Tfam** | **Tfb1m** | **Tfb2m** |
| --- | --- | --- | --- | --- |
| **1** | **Human** | NM_003201 | NM_016020 | NM_022366 |
| **2** | **Rat** | NM_031326.1 | NM_181474.2 | NM_001008293.1 |
| **3** | **Mouse** | NM_009360 | NM_146074 | NM_008249 |
| **4** | **Chimp** | NM_001194942.1 | NM_001362392.1 | ENSPTRT00000004012.2 |
| **5** | **Orangutan** | NM_001131295.1 | NM_001133415.1 | -NA- |
| **6** | **Gorilla** | NM_001131295 | NM_001292158 | ENSGGOT00000001101.1 |
| **7** | **Rhesus Monkey** | NM_001194942 | NM_001261482.1 | ENSMMUT00000032256.3 |
| **8** | **Green Monkey** | NM_001130211 | NM_001362392 | ENCSAT00000007204.1 |
| **9** | **Gibbon** | ENSNLET00000015683.2 | NM_ENSNLET00000037292.2 | NM_ENSNLET00000005278.3 |
| **10** | **Macaque** | ENSNFAT00000025894.1 | NM_001283474.1 | NM_001319608.1 |
| **11** | **Dog** | ENSCAFT00000038143.2 | NM_ENSCAFT00000000895.3 | -NA- |
| **12** | **Pig** | ENSSSCT00000046982.1 | NM_001128475.1 | ENSSSCT00000032587.2 |
| **13** | **Cow** | NM_001034016.2 | NM_001076896.2 | NM_001038127.2 |

**Table S3: Conditions for Western blotting**

| **S.no** | **Antibody** | **Blocking** | **Primary Antibody** | **Secondary antibody** |
| --- | --- | --- | --- | --- |
| **1** | Tfam | Overnight at 4˚C with 1% PVP | 1:7500  1 hour at RT | 1:5000  (Rabbit IgG) |
| **2** | Tfb1m | Overnight at 4˚C with 1% PVP | 1:5000  1 hour at RT | 1:3000  (Rabbit IgG) |
| **3** | Tfb2m | Overnight at 4˚C with 1% PVP | 1:5000  1 hour at RT | 1:3000  (Goat IgG) |
| **4** | HIF1α | 2 hours at RT with 5% skim milk | 1:2500  Overnight at 4˚C | 1:3000  (Mouse IgG) |
| **5** | PGC1α | 1 hour at RT with 5% skim milk | 1:3000  Overnight at 4˚C | 1:3000  (Rabbit IgG) |
| **6** | OXPHOS cocktail | 1 hour at RT with 5% skim milk | 1:400  Overnight at 4˚C | 1:4000  (Rabbit IgG) |
| **7** | Vinculin | 1 hour at RT with 3% BSA | 1:10000  Overnight at 4˚C | 1:5000  (Mouse IgG) |
